# Self-Organized Vascularized Cardiac Microtissues Derived from Human iPS Cells Promote Myocardial Repair through Functional Host–Graft Vascular Integration

**DOI:** 10.64898/2026.02.04.703917

**Authors:** Keisuke Hakamada, Kozue Murata, Wusiman Maihemuti, Kenji Minatoya, Hidetoshi Masumoto

**Author notes:** **Corresponding author:** Hidetoshi Masumoto, MD, PhD Department of Cardiovascular Surgery, Graduate School of Medicine, Kyoto University 54 Shogoin-kawahara-cho, Sakyo-ku, Kyoto, 606-8507, Japan, /.

## Abstract

**Objectives:** Cardiac regenerative therapy using human induced pluripotent stem cell (hiPSC)-derived tissues and organoids holds great promise for treating heart diseases. Successful clinical translation requires biomimetic cardiac tissues that not only recapitulate native myocardial architecture but also actively integrate with host vasculature. We aimed to engineer self-organized, vascularized cardiac microtissues (VCMs) and evaluate their therapeutic and regenerative potential in a rat model of myocardial infarction (MI).

**Methods:** VCMs composed of hiPSC-derived cardiomyocytes, vascular endothelial cells, and vascular mural cells were cultured under dynamic conditions to promote self-organization and prevascular network formation. One week after MI induction by coronary artery ligation in athymic immunodeficient rats, VCMs were transplanted onto the infarcted myocardium. Cardiac function was assessed by echocardiography and magnetic resonance imaging. Three-dimensional host–graft vascular architecture was visualized by light-sheet fluorescence microscopy following tissue clearing, and functional perfusion was evaluated by intravenous DyLight 488–conjugated lectin injection via host systemic circulation prior to tissue harvest.

**Results:** VCM transplantation significantly improved cardiac function and reduced infarct size compared with controls. Histological analyses demonstrated enhanced graft survival and neovascularization. Three-dimensional imaging revealed human-derived self-organized vascular networks within engrafted VCMs. Lectin perfusion confirmed functionally perfused, reciprocal host–graft vascular integration, including extension of graft-derived vessels into host myocardium, accompanied by myocardial regeneration. Early graft engraftment was significantly higher in the VCM group than in non-prevascularized controls.

**Conclusions:** Self-organized prevascularization of hiPSC-derived cardiac microtissues enable active host–graft vascular integration through functional vascular networks, thereby enhancing myocardial regeneration and therapeutic efficacy. This strategy represents an advanced approach for cardiac regenerative medicine.

**Summary:** This study aimed to develop self-organized, vascularized cardiac microtissues (VCMs) derived from human induced pluripotent stem cells (hiPSCs) and to evaluate their myocardial regenerative potential in a rat model of myocardial infarction (MI). VCMs were engineered from hiPSC-derived cardiomyocytes, endothelial cells, and vascular mural cells and cultured under dynamic conditions to enable self-organization and prevascular network formation. One week after MI induction, VCMs were transplanted onto the infarcted myocardium. Cardiac function was evaluated using echocardiography and magnetic resonance imaging. Light-sheet fluorescence microscopy combined with tissue clearing was used to visualize three-dimensional vascular architecture and host–graft integration, while lectin perfusion analysis assessed functional blood flow. VCM transplantation significantly improved cardiac function, increased early graft engraftment, and enhanced neovascularization. Importantly, self-organized human-derived vascular networks within the VCMs actively integrated with the host vasculature, forming functional, perfused host–graft vascular connections. These findings indicate that prevascularized VCMs do not merely survive after transplantation but actively promote vascular integration and myocardial regeneration through functional vascular networks. Together, these results demonstrate that self-organized vascularization markedly enhances graft integration, survival, and therapeutic efficacy, underscoring the clinical potential of VCM-based strategies for cardiac regenerative therapy.

## Introduction

The aging of the population is expected to lead to a rise in the number of patients experiencing severe heart failure^1^. Following hospitalization for heart failure, the overall survival rate over a 5-year period is a discouraging 50%^2,3^. Current treatment options for severe or refractory heart failure encompass cardiac transplantation and mechanical cardiac support (MCS) with the aim of extending life and enhancing the quality of life^2–4^. However, the supply of donor hearts for transplantation is quite restricted, and the long-term survival outlook for patients relying on MCS remains uncertain^4,5^. Consequently, there is an imperative need to explore new therapies for severe heart failure in order to broaden the scope of treatment for a larger number of heart patients.

Cardiac regenerative therapy is recognized as a promising approach for treating severe heart failure^6^. Clinical trials using somatic stem cells, such as bone marrow-derived cells and skeletal myoblasts, have shown some beneficial effects^7–9^. However, these trials could not demonstrate sufficient therapeutic efficacy as a standardized treatment, possibly due to lower differentiation into heart-constituent cells. Pluripotent stem cells [embryonic stem cells (ESCs), induced pluripotent stem cells (iPSCs)] are considered as ideal cell sources in context of higher efficiency of the differentiation into heart-constituent cells and have shown significant therapeutic efficacy in several pre-clinical studies^10–14^ which motivated to the implementation of clinical trials^15–17^. However, the therapeutic effect of pluripotent stem cells is mainly paracrine-mediated, involving pro-angiogenesis, anti-apoptosis, anti-inflammation, and anti-fibrosis. To address the limitations of a therapeutic mechanism primarily based on indirect paracrine factors, some researchers have attempted to improve cardiac function through intramyocardial transplantation with human iPSC-derived cardiomyocytes (CMs) in pre-clinical and clinical paradigm aiming for remuscularization of the heart^13,17,18^. However, these transplanted cells consist exclusively of CMs and lack vascular structure, preventing them from immediately functioning as part of the cardiac structure after transplantation.

In the current landscape of iPSC-based regenerative medicine, the transition from a single-cell transplantation approach to “organoid” transplantation is gaining momentum in various organs, potentially enhancing post-transplant tissue function^19,20^. However, endeavors to create tissue that mimics the native heart tissue, consisting of various heart cells, are still in progress within the field of cardiovascular regenerative medicine. We have been developing self-organized biomimetic cardiac tissues derived from human iPSCs that incorporate intrinsic vascular networks through dynamic rocking culture^21^. Unlike conventionally engineered constructs that rely primarily on externally imposed architecture, these tissues undergo active self-organization, allowing multiple cardiac cell types to autonomously assemble into higher-order structures that more closely resemble native myocardium. Through this process, the resulting tissues recapitulate not only the cellular composition but also the spatial organization and vascular integration characteristic of the human heart. As such, these self-organized tissues can be regarded as “cardiac organoids”, representing a distinct and biologically driven approach to cardiac tissue engineering. This strategy may introduce a new paradigm in cardiac regenerative medicine, in which active tissue self-organization may endow transplanted grafts with more physiologically relevant structure and function, thereby enhancing their potential to restore damaged myocardium in diseased hearts.

In this study, we introduce a novel approach utilizing “human iPSC-derived vascularized cardiac microtissues (VCMs)” in a rat model of myocardial infarction. What distinguishes our new graft is its self-organized robust vascular network, which is formed by human vascular cells from the in vitro culture stage. The presence of a well-organized vascular network in the graft holds the potential for establishing a functional vascular network capable of blood perfusion, thereby enhancing tissue regeneration capabilities. This development serves as a steppingstone toward creating ideal grafts that can directly contribute to the restoration of damaged myocardium.

## Methods

### Maintenance of human iPSCs and differentiation of cardiovascular cells

We employed two distinct human iPSC lines in this study: the 201B6 line, generated using four reprogramming factors (Oct3/4, Sox2, Klf4, and c-Myc)^22^, and the FfI01s04 line, established with a six-factor episomal plasmid vector (Oct3/4, Sox2, Klf4, L-Myc, LIN28, and Glis1)^23^, which is a human leukocyte antigen (HLA)-homozygous iPSC line developed at the Center for iPS Cell Research and Application (CiRA), Kyoto University, Kyoto, Japan, and distributed in accordance with informed consent from a healthy donor and approvals from the Institutional Review Boards of both RIKEN (RIKEN-K1-2023-001) and Kyoto University (G1025).

The maintenance of human iPSCs and their differentiation into cardiovascular cell lineages were performed following protocols from our previous studies, with some modifications^24^. Briefly, iPSCs were cultured and expanded in StemFit AK02N medium (AJINOMOTO, Tokyo, Japan). Upon reaching confluence, cells were dissociated using TrypLE Select (Thermo Fisher Scientific, Waltham, MA, USA), resuspended in a 1:1 solution of 0.5 mM ethylenediaminetetraacetic acid (EDTA) in phosphate-buffered saline (PBS), and passaged as single cells at a density of 5,000–8,000 cells/cm² every 5–7 days in AK02N medium supplemented with iMatrix-511 silk (0.125 μg/cm², FUJIFILM Wako Pure Chemical Corp., Osaka, Japan) and ROCK inhibitor Y-27632 (10 μM, FUJIFILM Wako). Penicillin–streptomycin (Thermo Fisher Scientific, 10,000 U/mL) was added as needed at a 1:100 dilution.

For CMs / endothelial cells (ECs) differentiation, single iPSCs were seeded on Matrigel-coated plates (1:60 dilution) at a density of 300,000–400,000 cells/cm² in AK02N medium containing 10 μM Y-27632. After reaching confluence, cells were overlaid with Matrigel (1:60 dilution in AK02N) one day prior to differentiation induction. Differentiation was initiated (day 0; d0) by replacing the medium with RPMI + B27 medium [RPMI 1640 (Thermo Fisher), 2 mM L-glutamine (Thermo Fisher), and 1× B27 supplement without insulin (Thermo Fisher)], supplemented with 100 ng/mL Activin A (R&D Systems, Minneapolis, MN, USA) and 5 μM CHIR99021 (Tocris Bioscience, Bristol, UK) for 24 hours. On day 1 (d1), the medium was supplemented with 10 ng/mL bone morphogenetic protein 4 (BMP4; R&D) and 10 ng/mL basic fibroblast growth factor (bFGF), and cultured for 4 days without medium change. On day 5 (d5), the medium was replaced with RPMI + B27 medium containing 50 ng/mL vascular endothelial growth factor (VEGF)_165_ (FUJIFILM Wako), and small molecules IWP4 (2.5 μM, Stemgent, Cambridge, MA, USA) and XAV939 (5 μM, Merck, Kenilworth, NJ, USA) were added from d5 to d7. Afterward, the culture medium was refreshed every other day using RPMI + B27 medium supplemented with VEGF—50 ng/mL for the 201B6 line and 12.5 ng/mL for the FFI01s04 line. Spontaneously beating CMs typically emerged between days 11 and 15.

To control the proportion of vascular mural cells (MCs) suitable for forming cell sheets, a portion of the differentiation culture was directed toward MC differentiation. The procedure up to day 3 was nearly identical to the CMs / ECs differentiation method described above, except that CHIR99021 was not used on day 0. After day 3, the medium was replaced with RPMI + FBS medium [RPMI1640 with 2 mM L-glutamine and 10% fetal bovine serum (FBS)] and changed every other day thereafter.

To prepare non-vascularized cardiac microtissues (NVCMs), we employed a differentiation protocol that specifically induces CMs without ECs. This protocol was largely similar to the one used for CMs / ECs induction described above, but with two key modifications: from day 5 onward, VEGF was omitted from the culture medium, and the 1× B27 supplement without insulin in RPMI + B27 medium was replaced with a 1× B27 supplement containing insulin (Thermo Fisher). During days 5–7, IWP4 and XAV939 were used as in the standard CMs / ECs induction protocol.

### Preparation of human iPSC-derived cardiac tissues

Following differentiation (days 13–15), cells were dissociated using Accumax (Innovative Cell Technologies, San Diego, CA, USA), and analyzed by flow cytometry. The dissociated cells were then mixed in accordance with the results of flow cytometry, and seeded onto FBS-coated 12-well UpCell plates (CellSeed, Tokyo, Japan) at a density of 1,500,000 to 4,000,000 cells per well in 2 mL of attachment medium (AM), consisting of alpha minimum essential medium (αMEM; Thermo Fisher) supplemented with 10% fetal bovine serum (FBS) and 5 × 10⁻⁵ M 2-mercaptoethanol, along with 25–50 ng/mL VEGF and 10 μM Y-27632. After two days of culture, VEGF (25–50 ng/mL) was added to the medium again. Two days later, the culture plates were transferred to room temperature. Within 15–30 minutes, cells detached spontaneously and floated in the medium as monolayer cardiac tissue sheets (CTSs). These detached CTSs were collected, transferred onto Matrigel-coated dishes, to promote reattachment, and incubated in AM containing 50 ng/mL Y-27632 for 24 hours.

For the preparation of VCMs and NVCMs, cardiac microtissues were cultured under dynamic rocking conditions, as described previously^24^. Specifically, the reattached tissues were placed on a Compact Digital Rocker (Thermo Fisher, #88880019) set to 60 rpm with a 13-degree tilt for 14 days. The culture medium was refreshed with AM every four days. As a control group for the VCMs, CTSs were prepared by culturing under static conditions for the same duration instead of dynamic culture.

### Histological analysis for human iPSC-derived cardiac tissues

Cardiac tissues were fixed in 4% PFA and embedded in paraffin. Sections were prepared and stained with hematoxylin-eosin and Sirius Red staining (FUJIFILM Wako Pure Chemicals). For immunohistochemical staining, paraffin sections were stained first with antibodies for cTnT (#MS-295-P0; Thermo Fisher) (1:100). They were incubated with biotinylated second antibody (1:300) and Avidin-biotin-peroxidase complex (ABC) (ABC-Elite, Vector Laboratories, Newark, CA, USA) (1:100). Coloring reaction was carried out with DAB and nuclei were counterstained with hematoxylin.

For immunofluorescence analysis, cardiac tissues were stained with a polyclonal rabbit anti-cardiac troponin-T antibody (#ab45932; Abcam, Cambridge, UK) (1:250), CD31 (monoclonal mouse IgG1, clone 9G1) (R&D) (1:250) with DAPI (4’,6-diamidino-2-phenylindole) (Thermo Fisher) (1:1000). Anti-rabbit Alexa 546 (Thermo Fisher) and anti-mouse Alexa Fluor 488 (Thermo Fisher) were used as secondary antibodies. The tissues were photographed with an all-in-one fluorescence microscopic system, BZ-X800E (Keyence, Osaka, Japan) Combined Z-stack and sectioning functions. All results were confirmed with >2 repetitive independent experiments.

### MUSCLEMOTION analysis (video-based method to evaluate tissue contractility)

MUSCLEMOTION is an adaptable open-source tool that utilizes video analysis to assess contractile function^25^. Following the guidelines of the provider, we implemented the software by installing it as a plug-in for ImageJ. The motion amplitude was measured for analysis.

### RNA sequencing (RNA-seq)

The VCMs and CTSs were dissociated using Accumax (Innovative Cell Technologies) to collect single dissociated cells. Total RNA was extracted using the RNeasy Mini Kit (QIAGEN, Hilden, Germany). Subsequently, reverse transcription to cDNA was performed using the QuantiTect Reverse Transcription Kit (QIAGEN). The cDNA library was prepared using the Next GEM Single Cell 39 Gel Bead Kit v3.1 (1000129), Chromium Next GEM Chip G Single Cell Kit v3 (PN-1000127), Next GEM Single Cell 39 GME Kit v3.1 (1000130), Next GEM Single Cell 39 Library Kit v3.1 (1000158), and i7 Multiplex Kit (PN-120262) (10x Genomics, Pleasanton, CA, USA) according to the instructions. Then the cDNA library was run on NextSeq 500 and HiSeq 4000 by Illumina (San Diego, CA, USA). For the quantification of gene expression, an analysis pipeline of ENCODE project (2.3.4) was used (https://www.encodeproject.org/pipelines/ENCPL002LPE/). GRCh38 ENSEMBL release 109 was used for the reference sequence. GRCh38 GENCODE release 43 was used for the gene definition. The genes in which Log_2_FC > Log_2_(1.5) were identified as “upregulated”, and the genes in which Log2FC < -Log2(1.5) were identified as “downregulated” in VCM group. Differential gene expressions were visualized using M-A plots generated in RStudio (version 4.4.0). Gene Ontology (GO) enrichment and network analysis were performed using ShinyGO (version 0.82) http://bioinformatics.sdstate.edu/go/.

### Animals

We purchased male immunodeficient athymic nude rats (F344/NJcl-rnu/rnu) from CLEA Japan Inc. (Tokyo, Japan) and housed them under controlled environmental conditions. All animal experimental procedures and the publication of related data were approved by the Animal Experimentation Committee of Kyoto University (Approval No. Med Kyo 22185). All experiments were conducted in accordance with the *Guidelines for Animal Experiments of Kyoto University*, which are consistent with Japanese law and the *Guide for the Care and Use of Laboratory Animals*.

### Induction of Myocardial infarction

The method for establishing the myocardial infarction (MI) rat model has been described previously^26–28^. All procedures were performed under general anesthesia, maintained with 2% isoflurane, and included respiratory support using a ventilator. Following anesthetization and endotracheal intubation, a left anterolateral thoracotomy was performed to expose the heart. MI was induced by permanent ligation of the left anterior descending (LAD) coronary artery.

### Transplantation of human iPSC-derived cardiac tissues

On day 7 following MI induction, human iPSC-derived cardiac tissue sheets were transplanted onto the anterior surface of the heart, as previously described^28^. The sheets were manually spread to cover the entire infarct and border zones and adhered to the heart surface without the need for sutures. In the Sham group, re-thoracotomy was performed in the same manner, and the chest was closed without transplantation.

### Evaluation of cardiac function with echocardiography and magnetic resonance imaging *(MRI)*

Echocardiographic assessments were conducted using the Vivid 7 system (GE Healthcare, Chicago, IL, USA) at baseline (prior to LAD ligation; pre-MI), on the day of transplantation (7 days post-MI; pre-Tx), and at 2-, 4-, and 12-weeks following transplantation. Parameters recorded included left ventricular end-diastolic diameter (LVDd), left ventricular ejection fraction (LVEF), and akinetic length.

At the end of the observation period, cardiac function was further evaluated using cardiac MRI with a 7-T BioSpec 70/20 USR system (Bruker Biospin, Ettlingen, Germany) under general anesthesia with 2% isoflurane (Pfizer). Left ventricular end-diastolic volume (LVEDV) and end-systolic volume (LVESV) were quantified using ImageJ software^29^, and LVEF was calculated as follows: LVEF (%) = 100 × (LVEDV − LVESV) / LVEDV.

### Histological analysis for rat hearts

At the end of the observation period, the animals were euthanized, and heart tissue samples were harvested and fixed in 4% paraformaldehyde (PFA). Paraffin-embedded tissue blocks were prepared, and serial sections (5 μm thickness) were collected at 50 μm intervals along the short axis of the heart from the site of LAD artery ligation. These sections were subjected to Sirius Red staining (FUJIFILM Wako Pure Chemicals) for the quantification of infarct size as the ratio of the infarcted endocardial length to the total inner circumferential length based on Sirius Red staining, using ImageJ software.

Immunostaining for cardiac isoform of cardiac troponin-T (cTnT) was performed using a polyclonal rabbit anti-cardiac troponin-T antibody (#ab45932; Abcam, Cambridge, UK) (1:1000). Visualization was carried out using the conventional HRP/DAB method with Avidin-biotin-peroxidase complex (ABC-Elite, Vector Laboratories) (1:100). Immunohistochemical fluorescence staining for STEM121 and von Willebrand factor (vWF) were conducted, using a mouse monoclonal STEM121 antibody specific for human cytoplasmic marker (#Y40410; Takara Bio Inc., Kusatsu, Japan) (1:400) and a rabbit polyclonal anti-vWF antibody (#ab6994; Abcam) (1:1000) was used as the primary antibody, followed by an Alexa Fluor 546-conjugated anti-mouse secondary antibody (Thermo Fisher; 1:400) and an Alexa Fluor 488-conjugated anti-rabbit secondary antibody (Thermo Fisher; 1:400).

All results were confirmed with >2 repetitive independent experiments. Histological images were acquired and analyzed using an all-in-one fluorescence microscope (Biorevo BZ-X810; Keyence, Osaka, Japan).

### Light sheet fluorescence microscopy

At 4 weeks post-transplantation, rats were sacrificed to evaluate the presence of human cells and vascular structures within the engraftment area using light sheet fluorescence microscopy (LSFM). To enable three-dimensional visualization, tissue clearing was performed using the CUBIC (Clear, Unobstructed Brain Imaging Cocktails and Computational analysis) protocol^30^. Briefly, excised heart tissues were first fixed in 4% PFA and washed in PBS for 24 hrs. The samples were then sequentially processed as follows: incubation in Tissue Clearing Reagent CUBIC-L (Tokyo Chemical Industry, Tokyo, Japan) at 37 °C for 14 days; immunostaining with rabbit polyclonal anti-vWF antibody (Abcam; 1:1000) and mouse monoclonal STEM121 antibody (1:500) for 3 days; followed by secondary antibody incubation with Alexa Fluor 488-conjugated anti-rabbit and Alexa Fluor 546-conjugated anti-mouse antibodies (Thermo Fisher Scientific; 1:400 each) for 3 days. Finally, samples were immersed in Tissue Clearing Reagent CUBIC-R (Tokyo Chemical Industry) for 1 day before imaging. All results were confirmed with >2 repetitive independent experiments.

### Lectin perfusion analysis

We labeled human iPSC-derived cardiac tissues with Hoechst 33342 to identify grafted cells after transplantation. The tissues were transplanted onto the epicardial surface of the heart one week after MI induction. One week after transplantation, rats were intravenously injected with 1.5 mL of 1 mg/mL DyLight 488-conjugated tomato (Lycopersicon esculentum) lectin (Vector Labs, Burlingame, CA, USA) via the inferior vena cava, 15 minutes before sacrifice, to visualize perfused vasculature. Following heart excision, the tissue samples were fixed in 4% PFA and embedded in paraffin. Immunohistochemical and immunofluorescence staining for von vWF were performed as described above. All results were confirmed with >2 repetitive independent experiments.

### Statistical analysis

All results are expressed as mean ± standard deviation (SD). Statistical analyses were performed using non-parametric methods. For comparisons between two groups, the Mann–Whitney U test was applied using GraphPad Prism 10 for Windows (version 5.03, GraphPad Software, Inc., San Diego, CA, USA). For comparisons involving more than two groups, non-parametric one-way analysis of variance by ranks was conducted, followed by post hoc analysis using the Steel–Dwass test for multiple comparisons.

### Data availability

Data supporting the findings of this study are available within the paper and its Supplementary Information.

## Results

### Tissue structure and function of human iPSC-derived VCMs

We first created self-organized VCMs for transplantation and evaluated their structures and tissue functions (Fig. 1A). The differentiation protocol for cardiovascular cell lines is shown in Fig. 1B and was used to generate thin cardiac tissue sheets (CTSs). These CTSs consisted of 31.8 ± 10.6% cardiac troponin T (cTnT)-positive cardiomyocytes (CMs), 6.96 ± 6.33% vascular endothelial cadherin (VE-cadherin)-positive vascular endothelial cells (ECs), and 47.6 ± 10.4% platelet-derived growth factor receptor-beta (PDGFRβ)-positive vascular mural cells (MCs) (n = 7) (Fig. 1C). We generated VCMs by applying dynamic rocking culture to the CTSs. As a control group, we simultaneously prepared CTSs without self-organized vascular network formation that were maintained under static culture conditions for the same duration as the dynamic culture.

**Figure 1:**
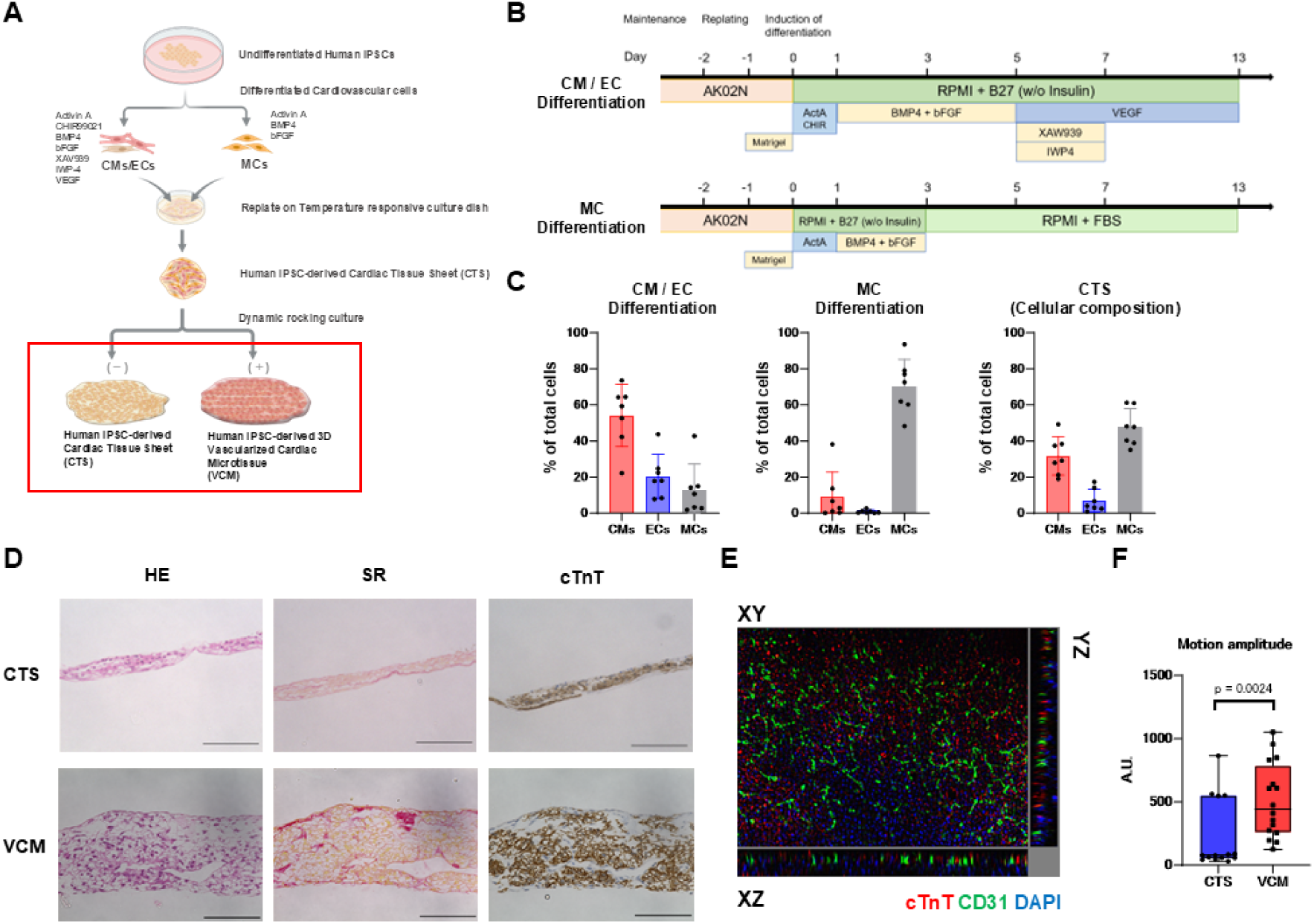
In vitro comparison between CTSs and VCMs. (A) Schematic diagram of human iPSC-derived 3D vascularized cardiac microtissues formation. The components highlighted with red squares (VCM and CTS) were used for comparative analysis in the subsequent experiments (excluding Figure 6). (B) Schematic representation of differentiation of cardiovascular cell lines Upper: cardiomyocytes [CMs], endothelial cells [ECs]; Lower: vascular mural cells [MCs]. (C) Cellular components after differentiation of cardiovascular cells derived from human iPSCs (n = 7). CMs, cardiomyocytes; ECs, vascular endothelial cells; MCs, vascular mural cells. (D) Representative histological evaluations of cardiac tissue sheet and vascularized 3D cardiac microtissues derived from human iPSCs. Upper: hiPSC-derived cardiac tissue sheet (CTS) Lower: hiPSC-derived 3D vascularized cardiac microtissues (VCM) Left: haematoxylin–eosin (H-E) staining; Middle: Sirius red (SR) staining; Right: cardiac isoform of troponin-T (cTnT) immunostaining. (Scale bar 100 *μ* m.) (E)) Immunofluorescence analysis (IFA) of VCM for cTnT (red; CMs), CD31 (green; ECs), and 4’,6-diamidino-2-phenylindole (DAPI) (blue; cell nuclei) after 14 days of rocking culture. (F) MUSLEMOTION analysis of CTS and VCM. (CTS: n = 14, VCM: n = 16). A.U., arbitrary unit.

Histological analysis revealed that VCMs were obviously thicker than CTSs. VCMs had an approximately 100-150 µm-thick cTnT-positive CM layer and were composed of around 15 cell layers in total. Sirius Red staining revealed that the VCMs formed layered collagen fiber structures, indicating the development of extracellular matrix (ECM), which is essential for tissue architecture and function (Fig. 1D). Fluorescent immunostaining of the VCMs indicated that they were composed of cTnT-positive CMs and vascular cells that were distributed among the myocardial layer. CD31-positive vascular endothelial cells formed an organized vascular network among the VCMs (Fig. 1E; Video 1). Results from the tissue contraction analysis using a video-based cardiac tissue motion assessment system^25^ showed that the motion amplitude of VCMs was significantly higher than that of CTSs (P = 0.0024) (Fig. 1F).

### Gene expression profiles of VCMs

In order to explore the impact of the structural maturation of the VCM on gene expression patterns, we conducted an RNA-seq analysis to compare gene expression profiles between the self-organized VCM and non-organized CTS groups. A total of 2,037 genes were found to be upregulated in VCM (Log_2_FC > log_2_(1.5)), whereas 633 genes were identified as downregulated (Log_2_FC < -log_2_(1.5)) (Fig. 2A). We employed an M-A plot analysis to visualize the distribution of differential gene expression. This plot clearly demonstrated increased expression of several genes, including DLK1, MGP, TFPI2, DCN, SERPINA1, and SNORD3D, in the VCM group (Fig. 2B). Subsequently, Gene Ontology (GO) enrichment analysis was performed on the 2,037 upregulated genes. The analysis revealed significant enrichment in pathways related to ECM organization, blood vessel morphogenesis, and tissue development. Notably, terms such as “vasculature development,” “angiogenesis,” and “tube morphogenesis” were prominently enriched, suggesting that the upregulated gene set is closely associated with endothelial cell identity and vascular function (Fig. 2C,D). These findings highlight the potential involvement of endothelial-related biological processes in the VCM group and underscore the importance of vascular components in the observed transcriptomic differences.

**Figure 2:**
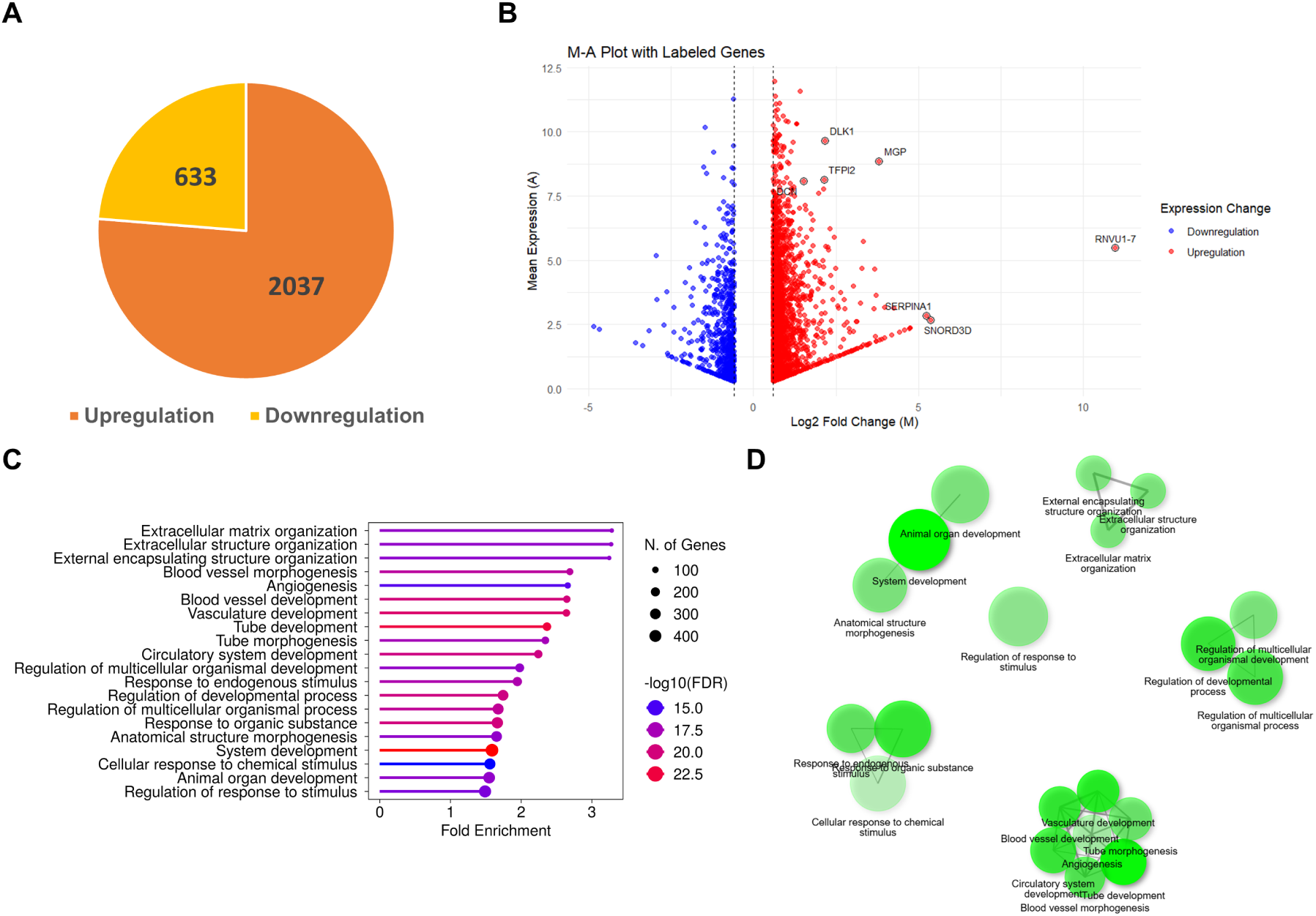
Transcriptomic analysis of VCMs. (A) RNA-seq analysis comparing gene expression between CTSs and VCMs revealed 2,037 genes upregulated (Log2FC > log2(1.5)) and 633 genes downregulated (Log2FC < –log2(1.5)) in the VCM group. (B) M–A plot showing differentially expressed genes. Notable genes with increased expression in VCMs include DLK1, MGP, TFPI2, DCN, SERPINA1, and SNORD3D. (C, D) Gene Ontology (GO) enrichment analysis of the upregulated genes in VCMs.

### Transplantation of VCMs ameliorated cardiac dysfunction after MI

We next performed the transplantation of self-organized VCMs into a rat model of MI to confirm their therapeutic potential. There were no significant differences among the three groups in echocardiographic parameters before transplantation (after MI induction). Echocardiogram at 4- and 12-weeks post-transplantation of VCMs showed a lower left ventricular diastolic dimension (LVDd), lower akinetic length, and higher left ventricular ejection fraction (LVEF) compared to the other groups (Sham vs CTS vs VCM, 4 weeks: LVDd, 0.73 ± 0.04 vs 0.71 ± 0.05 vs 0.65 ± 0.04 mm, P = 0.0155; LVEF, 49.5 ± 1.37 vs 60.9 ± 4.22 vs 76.4 ± 2.50 %, P < 0.0001; Akinetic length, 4.88 ± 0.81 vs 3.36 ± 0.85 vs 2.42 ± 0.55 mm, P = 0.0006; 12 weeks: LVDd, 7.5 ± 0.5 vs 6.7 ± 0.3 vs 6.0 ± 0.1 mm, P < 0.0001; LVEF, 47.5 ± 3.4 vs 60.5 ± 1.3 vs 76.1 ± 1.7 %, P < 0.0001; Akinetic length, 5.6 ± 0.7 vs 4.2 ± 0.2 vs 2.4 ± 0.2 mm, P < 0.0001) (Fig. 3A; Supplemental Fig. 1). Cardiac MRI at 4 and 12 weeks after transplantation revealed the highest LVEF in the VCM group (Sham vs CTS vs VCM, 4 weeks: 34.0 ± 3.7 vs 41.8 ± 2.3 vs 54.9 ± 2.5 %, P < 0.0001; 12 weeks: 34.6 ± 5.9 vs 40.4 ± 5.7 vs 56.5 ± 2.5 %, P < 0.0001) (Fig. 3B, C). These results indicate that VCM transplantation has superior therapeutic effects on a rat MI model.

**Figure 3:**
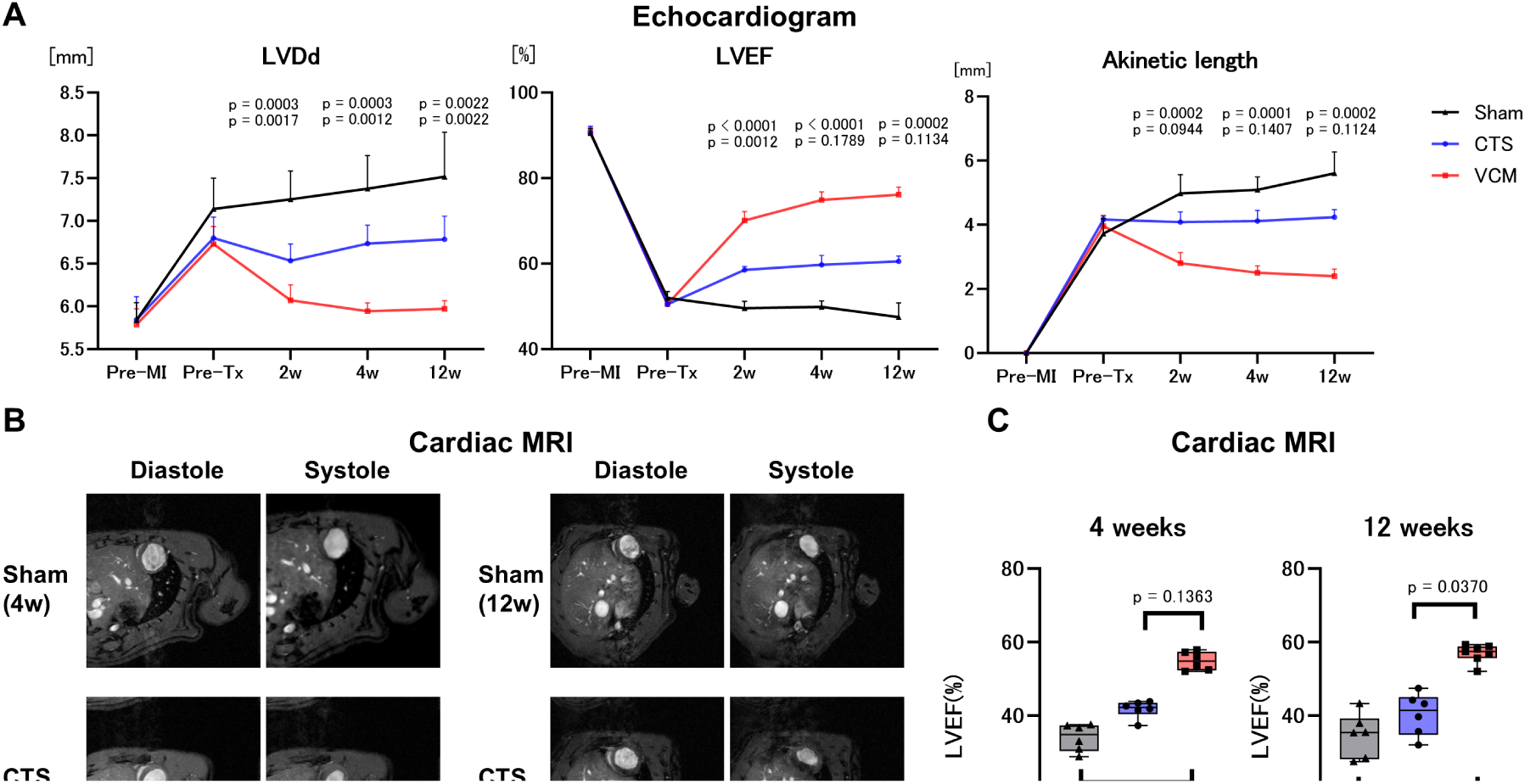
Cardiac function evaluation after VCMs and CTSs transplantation. (A) Echocardiography data over time post-transplantation: Left: left ventricular (LV) end-diastolic dimension (LVDd), Middle: LV ejection fraction (LVEF), and Right: LV Akinetic length. Upper p, VCM vs. Sham. Lower p, VCM vs. CTS. (B) Representative cardiac magnetic resonance imaging (MRI) images at 4 and 12 weeks after transplantation (C) Results for LVEF evaluated by cardiac MRI. Wilcoxon test, post hoc Steel–Dwass multiple-comparison test. Four-week observation: sham, n = 6; CTS, n = 6; VCM, n = 6; 12-week observation: sham, n = 6; CTS, n = 6; VCM, n = 7.

### Transplantation of VCMs restored human myocardium in the recipient rat heart after MI and attenuated left ventricular remodeling

Next, to evaluate the myocardial restoration after VCM transplantation, we measured the positive area of STEM121 immunostaining, a specific marker for human cytoplasmic cells. At 4- and 12-weeks post-transplantation, the VCM group showed a significantly larger engraftment area of human cells compared to the CTS group (CTS vs VCM, 4 weeks: 0.1 ± 0.07 vs 1.9 ± 1.4 mm², P = 0.0079; 12 weeks: 0 vs 0.26 ± 0.13 mm², P = 0.0012) (Fig. 4A). Furthermore, we evaluated the left ventricular remodeling by Sirius Red staining. The VCM group exhibited significantly smaller scar extension (ratio of infarct endocardial length to total inner circumflex length: Sham vs CTS vs VCM, 4 weeks: 0.56 ± 0.04 vs 0.42 ± 0.05 vs 0.30 ± 0.05, P < 0.0001; 12 weeks: 0.55 ± 0.04 vs 0.49 ± 0.06 vs 0.27 ± 0.04, P < 0.0001) (Fig. 4B). These results indicate that the self-organized VCM transplantation has excellent capacity for myocardial restoration and simultaneously exerts suppressive effects on left ventricular remodeling.

**Figure 4:**
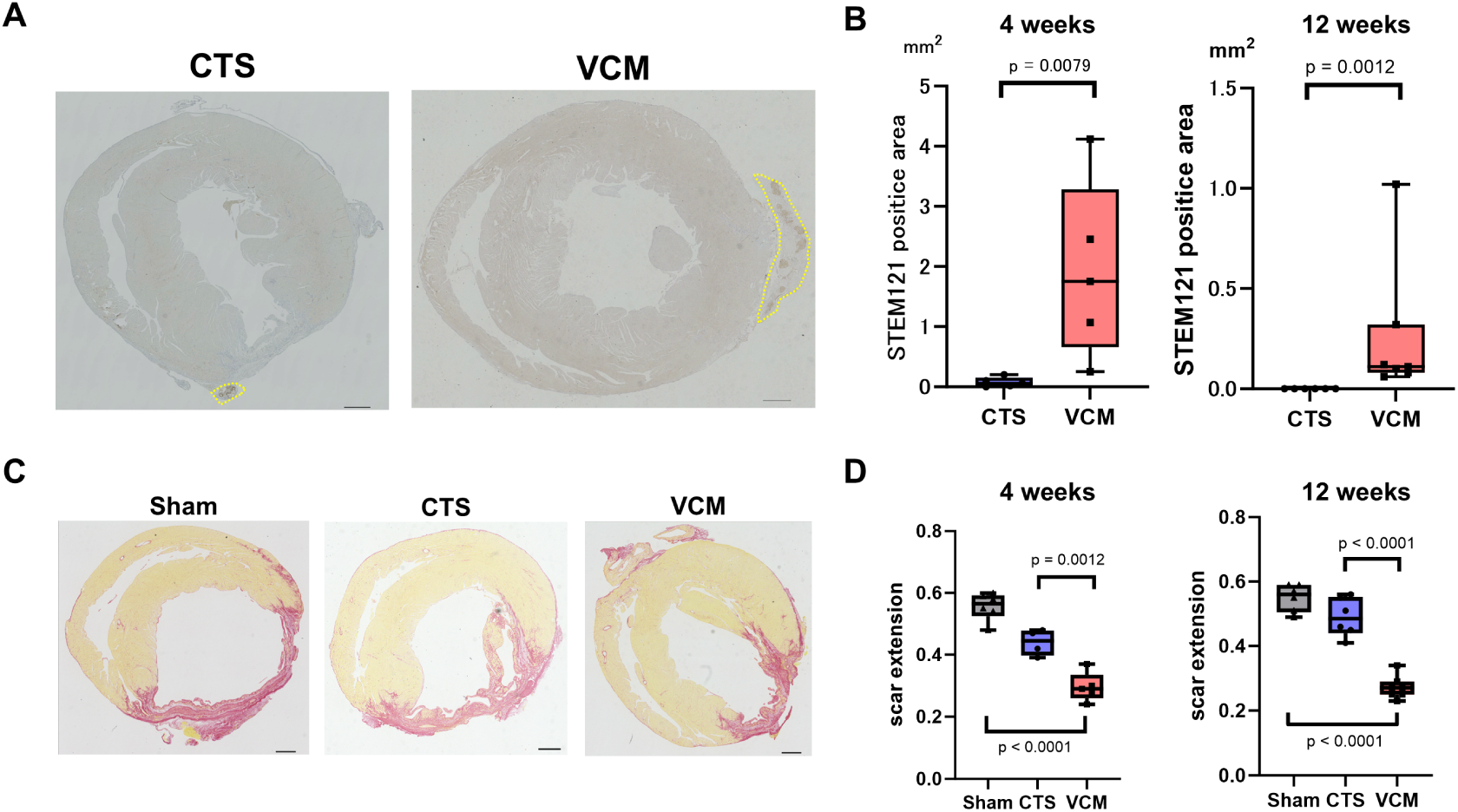
Histological analysis. (A) Representative immunostaining for STEM121 (brown / yellow dotted area) at 4 weeks after transplantation. Left and center panels, lower-magnification views. (Scale bars: 1000 μm). (B) Quantitative evaluation of the STEM121-positive area. P was calculated by Wilcoxon test. Four-week observation: CTS, n = 5; VCM, n = 5; 12-week observation: CTS, n = 6; VCM, n = 7. (C) Representative Sirius Red staining at 4 weeks after transplantation (Scale bars: 1000 μm). (D) Quantitative evaluations of scar extension (ratio of infarct endocardial length to total inner circumflex length). P was calculated by Wilcoxon/Kruskal–Wallis test, post hoc Steel–Dwass multiple-comparisons test. Four-week observation: sham, n = 6; CTS, n = 5; VCM, n = 5; 12-week observation: sham, n = 6; CTS, n = 6; VCM, n = 7.

### Transplantation of VCMs promoted neovascularization at the border zone of MI and functional vascular network formation composed of transplanted vascular cells among regenerated myocardium

Subsequently, we evaluated vascular density at the border zone of MI. VCM transplantation significantly increased the number of vWF-positive vasculatures in close proximity to the graft compared with the sham and CTS groups at 4 and 12weeks after transplantation [4 weeks, Sham vs CTS vs VCM: 1.7 ± 0.8vs 2.4 ± 1.1 vs8.1 ± 2.2 /mm^2^, *p* = 0.006 ; 12weeks 1.8 ± 0.8 vs 3.0 ± 0.9 vs 10.5 ± 2.2 / mm^2^, *p* < 0.0001] (Figure 5A,B).

**Figure 5:**
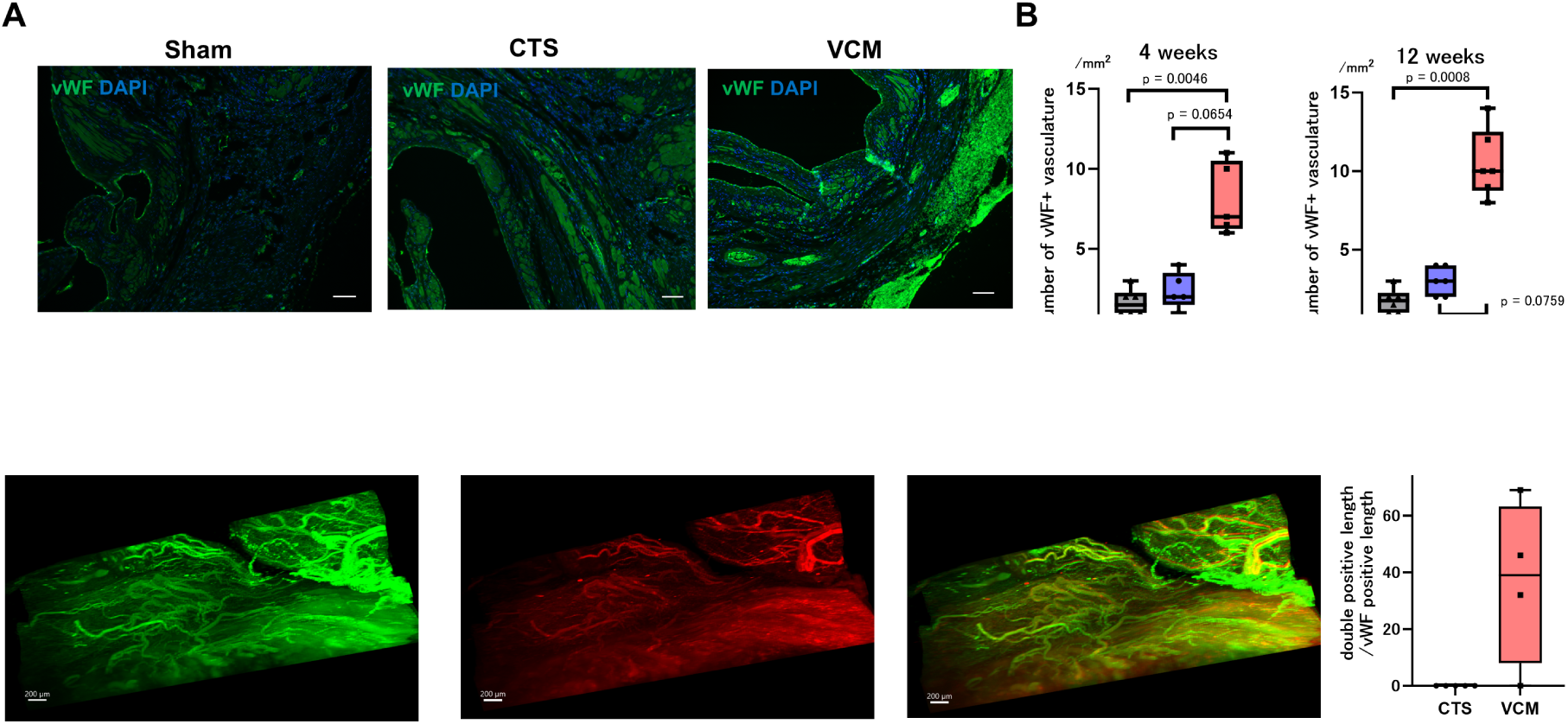
Evaluation of angiogenesis in the host MI border zone and vascular network formation within the engrafted tissue. (A) Representative immunofluorescence staining for von Willebrand factor (vWF) (green; endothelial cells [ECs]) and 4’,6-diamidino-2-phenylindole (DAPI) (blue; cell nuclei) at the border zone of MI at 4 weeks after transplantation. (Scale bars: 100 μm). (B) Quantitative evaluation of vascular density (/mm^2^) at 4 and 12 weeks after transplantation. P was calculated by Wilcoxon/Kruskal–Wallis test, post hoc Steel–Dwass multiple-comparison test. Four-week observation: sham, n = 6; CTS, n = 5; VCM, n = 5; 12-week observation: sham, n = 6; CTS, n = 6; VCM, n = 7. (C) Representative light sheet fluorescence microscopy (LSFM) image for von Willebrand factor (vWF) (green; endothelial cells [ECs]) and STEM121 (Red; human cells) at 4 weeks after VCM transplantation. Marked vascular network formation among the VCM grafts was observed at 4 weeks after transplantation. (Scale bars: 200 μm). (D) Quantitative evaluation of ratio of double positive length to vWF-positive length. P was calculated by Wilcoxon test. CTS, n = 5; VCM, n = 4.

To investigate the formation of the vascular network within the graft and its cellular origin, we utilized LSFM to evaluate human cells and vasculature among the grafts at 4 weeks after transplantation. LSFM analysis revealed the presence of vasculatures originating from human cells within the engrafted VCMs (Fig.5C). We confirmed that the ratio of the length of human vasculature [STEM121 and vWF double positive] to the length of total vasculature [vWF single positive] is significantly higher in the VCM group than that in the CTS group. [CTS vs VCM: 0 % vs 36.8 ± 14.7 %, *p* = 0.0476) (Fig. 5D). These findings provide evidence that transplanting tissues prevascularized with human blood vessels results in the structural establishment of a human-origin vascular network within the graft after transplantation.

### The functionality of the vascular network among the graft in terms of blood perfusion and its impact on early engraftment of transplanted tissues

To evaluate whether the vascular network within the engrafted tissue is functional and capable of supporting blood perfusion, we performed lectin perfusion analysis. Based on our previous findings, we concluded that non-organized CTS, which does not undergo dynamic training, exhibits low engraftment efficiency (Fig. 4A) and is therefore not suitable as a control group for evaluating the functional vasculature within engrafted tissue. As an alternative control for VCM, we prepared non-vascularized cardiac tissue (NVCM), which lacks endothelial cells but was subjected to the same dynamic training as VCM, resulting in a similar tissue thickness. Histological analysis prior to transplantation demonstrated that tissue thickness and the cardiomyocyte composition were comparable between VCM and NVCM (Figure 6A). We stained the nuclei of VCMs or NVCMs with Hoechst 33342 and transplanted them onto an immunodeficient rat model of MI. One week later, DyLight 488-conjugated lectin was injected via the inferior vena cava of the rats, and after sufficient perfusion throughout the heart, the rat hearts were harvested for histological sectioning. In this experimental model, transplanted grafts and host rat myocardium could be distinguished by Hoechst 33342 without the need for additional immunohistochemical staining. Furthermore, functional vascular networks with blood perfusion could be visualized regardless of whether they originated from the host or the graft (Fig. 6B; left). This lectin perfusion analysis demonstrated that there were functional vascular structures both with human nuclei (functional human vasculature; Fig. 6B-i) and without human nuclei (functional non-human vasculature; Fig. 6B-ii) within the VCMs, indicating that VCMs had a host-derived functional vascular network at 1 week after transplantation. Notably, vascular structures containing human nuclei were observed within Hoechst 33342-negative regions of the host rat myocardium (Fig. 6B-iii). This finding suggests that neovascularization from the engrafted VCMs had extended into the host rat heart tissue. It indicates that the regenerated human cardiac tissue resulting from self-organized VCM transplantation possesses the ability to extend its vasculature and integrate with the host vasculature, thereby forming an integrated vascular network that functionally connects the graft and host tissues. In contrast, transplantation of NVCMs did not result in the formation of a functional vascular network within the graft, unlike the VCM group.

**Figure 6:**
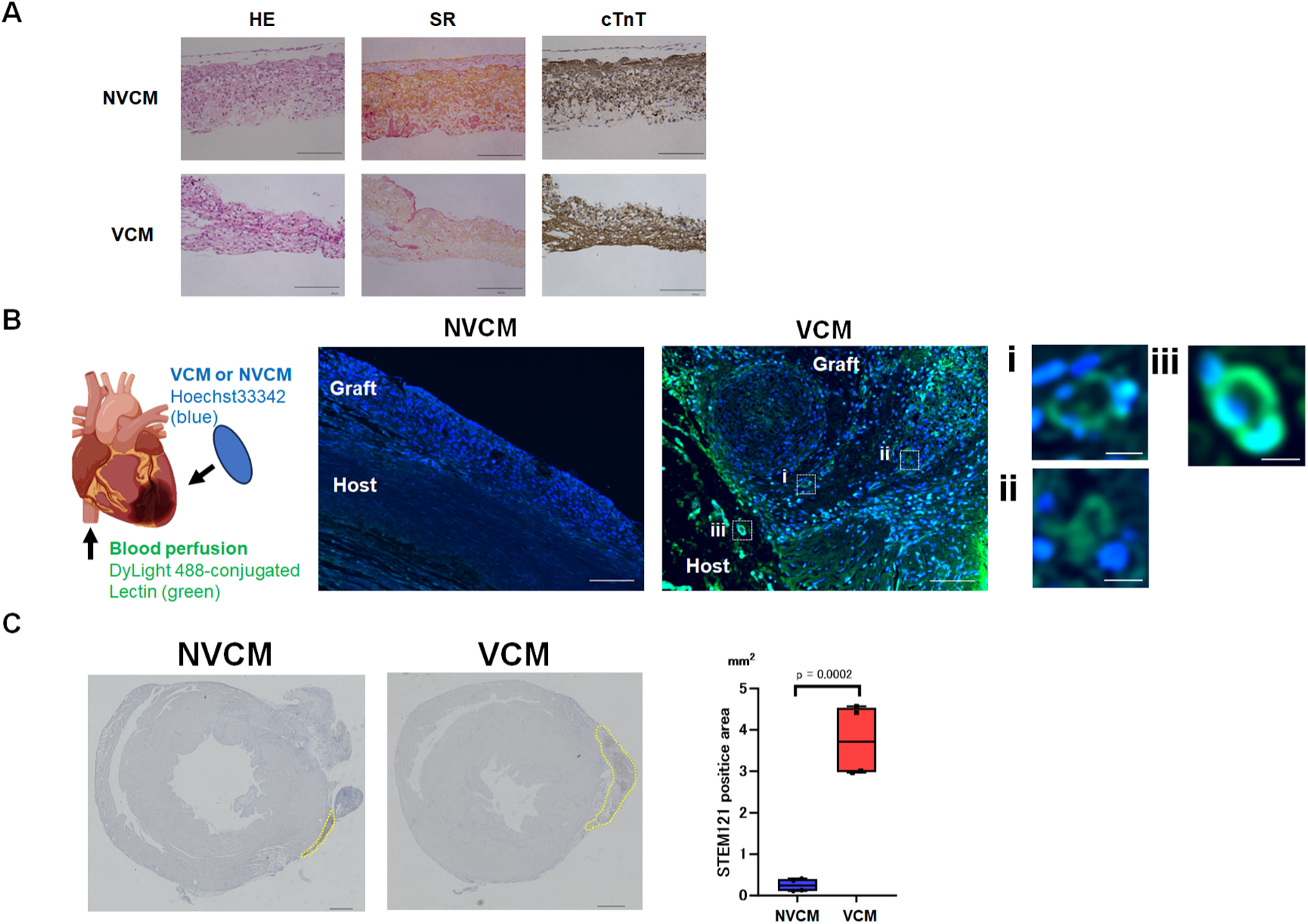
Lectin perfusion analysis. (A) Representative histological evaluations of NVCM and VCM. Left: Haematoxylin–Eosin (H-E) staining; Middle: Sirius Red (SR) staining; Right: cardiac isoform of troponin-T (cTnT) immunostaining (Scale bar 100 *μ* m). (B) Schematic diagram of lectin perfusion analysis (left). Representative immunofluorescence staining for DyLight 488-conjugated lectin (green; functional vasculature) and Hoechst33342 (blue; human nuclei) at 1 week after transplantation (Scale bars: 100 μm) (right). (i-iii) High magnification view (Scale bars: 10 μm). (C) Representative immunostaining images for vWF on sections used in the lectin perfusion analysis (left). Top: MI heart from a sham-operated control rat; Bottom: MI heart from a rat transplanted with VCM. Quantification of the classes of vasculatures (right). Sham, n = 2; NVCM, n = 4; VCM, n = 4. (D) Representative immunostaining for STEM121 (brown, yellow dotted area) at 1 week after transplantation (Scale bars: 1000 μm) (left). Quantification of the STEM121-positive area (right). P was calculated by Wilcoxon test. NVCM, n = 4; VCM, n = 4.

Finally, we investigated whether the presence of vasculature promotes early graft engraftment. One week after transplantation, histological analysis using STEM121 immunostaining revealed that the VCM group exhibited a significantly greater graft area compared to the NVCM group (CMT vs. VCM: 0.25 ± 0.15 vs. 3.74 ± 0.87 mm², P = 0.0002) (Fig. 6C). These findings suggest that prevascularization of cardiac tissues prior to transplantation enhances early engraftment efficiency and may contribute to improved therapeutic outcomes.

## Discussion

In this study, we successfully generated self-organized, vascularized cardiac microtissues derived from human iPSCs using dynamic rocking culture. Through this process, multiple cardiac cell types actively and autonomously assembled into higher-order tissue structures, resulting in the formation of robust intrinsic vascular networks already at the in vitro stage prior to transplantation. This capacity for active self-organization, rather than reliance on externally imposed architecture, represents a key distinguishing feature of our approach and may allow the engineered tissues to more closely recapitulate the structural and functional characteristics of native myocardium. Importantly, transplantation of these human iPSC-derived VCMs into a rat model of myocardial infarction demonstrated significant therapeutic efficacy and myocardial regeneration. Notably, this was accompanied by the formation of functional vascular networks composed of both human VCM-derived vessels and host-derived vessels, indicating successful and biologically integrated host–graft vascular coupling. Together, these findings suggest that self-organized tissue assembly enables grafts to achieve more physiologically relevant vascular integration, which may be critical for restoring myocardial structure and function in vivo.

Dynamic rocking culture is an approach designed to emulate the physiological in vivo environment. The mechanism of this system aims to facilitate the construction of a three-dimensional structure through the influence of biochemical factors and mechanical stimulation^31,32^. Vascular network formation within the grafts mediated by this dynamic culture system is considered to recapitulate self-organization during native organ development. Because such self-organizing processes are driven by cell–cell interactions and spatial patterning that give rise to in vivo–like structure and function^33^, this approach may enable the generation of tissue constructs with more physiologically relevant architecture and functionality than bioengineered cardiac tissue grafts fabricated using techniques such as three-dimensional bioprinting^34^. Furthermore, in contrast to existing grafts used in regenerative therapy, which typically require biomaterials like gelatin hydrogels to enhance thickness and support oxygen and nutrient supply^35^, VCMs exhibit thickness without the need for such biomaterials. This characteristic makes the graft more suitable for clinical use than existing grafts, as it may eliminate potential foreign body reactions upon transplantation. Additionally, VCMs feature vascular networks within the grafts. Angiogenesis, the process by which vascular cells sprout from pre-existing vessels to form new vasculature, is considered crucial for restoring tissue function and structure at sites of injury^36^. Our engineered VCMs possess pre-formed vascular networks prior to transplantation, indicating that this regenerative process is partially achieved before implantation. This prevascularization is likely to contribute positively to the establishment of functional, perfused vascular networks after transplantation. Furthermore, we demonstrated that the VCM actively induces angiogenesis toward the recipient heart, suggesting that the graft possesses intrinsic angiogenic potential that may play a key role in tissue repair and regeneration.

Through RNA-seq analysis, we were able to gain deeper insights into tissue architecture in the VCM group. In the GO enrichment analysis, GO terms related to ECM organization and structuring were significantly enriched, along with those associated with vascular formation and development. Furthermore, M-A plot analysis revealed upregulation of genes such as DCN, TFPI2, MGP, and SERPINA1 in the VCM group—genes known to regulate ECM and protease activity^37–40^. These are believed to contribute to tissue remodeling, control of fibrosis, and maintenance of structural integrity. Considering the crucial role of the ECM in tissue architecture and function, these findings are particularly compelling. Additionally, genes such as DLK1 and MGP are associated with developmental processes and cell differentiation^37,41^, suggesting a shift toward a more regenerative phenotype in the VCM group. Moreover, the non-coding RNAs RNVU1-7 and SNORD3D, which are involved in RNA splicing and modification, point to the involvement of epigenetic or post-transcriptional regulation of gene expression^42,43^. Taken together, the upregulation of these genes in the VCM group likely reflects enhanced ECM remodeling, activation of developmental signaling pathways, and increased regulatory RNA activity—possibly indicating changes in cellular identity and regenerative capacity.

We are now approaching an unprecedented super-aged society, particularly in developed countries, which has significantly accelerated the progress of regenerative medicine research. In the field of cardiovascular medicine, where the number and severity of heart failure cases are projected to increase, regenerative therapies using iPSC-derived organoids or cardiac tissues are expected to become realistic alternatives to heart transplantation. Among the available methods, transplanting cells in the form of sheets or patches offers the advantage of replicating the native structure of the heart, provided that the transplanted tissues are appropriately engineered as surrogate myocardial tissues^14,31,44^. This approach is based on the concept that the heart is an organic structure composed not only of CMs but also of various other cell types, including vascular endothelial cells, vascular smooth muscle cells, and fibroblasts. The self-organized VCMs used in this study exhibit a structure that closely aligns with this concept, resembling a “cardiac organoid.” In this study, we demonstrated that regenerated myocardial tissue equipped with a functional vascular network can be efficiently integrated into a diseased heart. This was achieved by transplanting tissues that already contained a pre-formed vascular network. Furthermore, transplantation of VCMs resulted in the recovery of cardiac function, and histological evaluation revealed suppression of left ventricular remodeling. Lectin perfusion analysis performed one week after transplantation demonstrated reciprocal formation of functional vascular networks, including graft-derived vasculature within the VCMs and their extension from the graft into the host myocardium. These findings suggest that the establishment of vascular networks may be a key strategy for efficiently restoring the function of damaged myocardium. The ultimate goal of transplantation using cardiac organoids is to regenerate myocardial tissue that actively contributes to the contractile function of the heart. Although our findings suggest favorable structural and functional integration, additional studies will help to further elucidate the contribution of regenerated myocardial tissue to overall cardiac contractility. Nevertheless, by further advancing the myocardial tissue regeneration strategy demonstrated in this study, we believe it may be possible to replenish a substantial portion of lost myocardium and complement the pumping function of the heart, potentially achieving therapeutic outcomes comparable to heart transplantation.

There are several limitations to this study. Firstly, there is the interspecific difference between rats and humans. This introduces the possibility that immune rejection may not be as relevant in humans. To address this, future experiments, such as allogenic transplantation, will be necessary. Secondly, the model of MI employed in our study is based on artificial coronary ligation in rats, which differs from the actual human pathophysiology characterized by atherosclerosis.

In conclusion, the transplantation of human iPSC-derived self-organized VCMs showed significant therapeutic potential in a rat model of MI associated with the development of a functional vascular network formed by transplanted human cells. This approach holds immense promise as an effective strategy for efficiently restoring the damaged myocardium in the future.

## Acknowledgements

We thank for Mrs. Kanako Takakura, Dr Kenta Terai, and Dr Michiyuki Matsuda (Kyoto University) for technical assistance with LSFM, and we thank the Center for Anatomical, Pathological and Forensic Medical Research, Kyoto University Graduate School of Medicine for preparing microscope slides and immunostaining. We thank Ms. Aiko Yoshioka (Kyoto University) for providing expert technical assistance.

## Declaration of generative AI and AI-assisted technologies in the writing process

During the preparation of this work the authors used ChatGPT in order to improve language and readability, with caution. After using this tool/service, the authors reviewed and edited the content as needed and take full responsibility for the content of the publication.

## Clinical Perspective

### What is New?

- We developed self-organized vascularized cardiac microtissues (VCMs) composed of hiPSC-derived cardiomyocytes, endothelial cells, and mural cells through dynamic culture.
- Transplantation of these VCMs into infarcted rat hearts resulted in improved cardiac function, enhanced cell engraftment, and host-graft vascular integration.
- Light-sheet fluorescence microscopy visualized 3D vascular networks formed by human and host-derived cells within the grafts.

### What are the Clinical Implications?

- Prevascularization of hiPSC-derived cardiac tissues may enhance engraftment efficiency and therapeutic efficacy in patients with ischemic heart disease.
- Self-organized VCMs offer a promising platform for future regenerative therapies aiming to restore cardiac function after myocardial infarction.
- This strategy could accelerate the clinical translation of stem cell-based cardiac tissue therapies by addressing key challenges in vascular integration.

